# Egg size does not universally predict embryonic resources and hatchling size across annual killifish species

**DOI:** 10.1101/2020.02.25.964411

**Authors:** Milan Vrtílek, Tom J. M. Van Dooren, Mégane Beaudard

## Abstract

Egg size has a crucial impact on the reproductive success of a mother and the performance of her offspring. It is therefore reasonable to employ egg size as a proxy for egg content when studying variation in offspring performance. Here, we tested species differences in allometries of several egg content parameters with egg area. We measured individual eggs in five species of annual killifish (Cyprinodontiformes), a group of fish where egg banks permit population survival over dry season. Apart from comparing allometric scaling exponents, amounts and compositions of egg components across the different species, we assessed the explanatory power of egg area for egg wet and dry weight and for hatchling size. We found notable species-specific allometries between egg area and the other egg parameters (egg dry weight and water content, elemental composition and triglyceride content). Across species, egg area predicted egg wet weight with highest power. Within species, coefficients of determination were largest in *Austrolebias elongatus*, a large piscivorous species with large eggs. Our study shows that systematically using egg area as a proxy of egg content between different species can ignore relevant species-specific differences and mask within-species variability in egg content.

## Introduction

Egg size is an ecologically and evolutionary important trait in egg-laying animal species (Bernardo, 1996; Fox and Czesak, 2000; Krist, 2011). In particular in species that provide little or no parental care, the egg and the resources it contains are the main route of transmission of non-genetic effects between a mother and its offspring. Parameters describing an egg thus do not only affect individual offspring performance, but also reflect female’s allocation strategy that eventually impacts her fitness (Moore et al., 2015; Morrongiello et al., 2012; Smith and Fretwell, 1974). Egg size usually provides a reasonable proxy for the other traits potentially important for offspring survival, development and growth; like the amounts of water, nutrients, or energy contained in the egg (Berg et al., 2001; Salze et al., 2005). Hence, we often observe long-term cascading effects of egg size on early-life offspring performance (Segers and Taborsky, 2011; Self et al., 2018; Semlitsch and Gibbons, 1990). Measuring egg size is straightforward and can be performed in a non-invasive manner, usually recorded as diameter or wet mass (Brooks et al., 1997; Krist, 2011; Räsänen et al., 2005). Using egg size as a proxy for other egg parameters seems, therefore, a sufficient way to assess, or control for female allocation among individual offspring.

Some studies have shed doubt on the generality of strong positive correlations between egg size and egg content (Leblanc et al., 2014; Moran and McAlister, 2009; Murry et al., 2008), showing that egg dry mass proves to be a more reliable predictor of nutrient and energy content (Murry et al., 2008). While we might measure maternal investment better by the means of egg dry weight, the problem is that destroying the egg by drying prevents any follow-up monitoring at the individual offspring level. Most egg parameters actually cannot be measured without destroying the embryo. Egg size therefore remains the first option to test for effects of maternal investment on offspring life history. To make valid predictions without sacrificing all eggs, we need good data on the relationship of egg size with other egg parameters from multiple species.

Annual killifish (Cyprinodontiformes) are small freshwater fish adapted to seasonally desiccating ponds where populations overcome dry periods in embryonic stages, thus creating egg banks in the pond soil (Cellerino et al., 2016; Furness, 2016). Adaptation to this temporary habitat evolved multiple times from non-annual ancestors (Furness et al., 2015; Helmstetter et al., 2016). A common feature of annual killifish embryos is their ability to arrest development (by entering diapause) and potentially reduce energy expenditure at three discernible morphological stages (Podrabsky et al., 2010; Wourms, 1972). Diapauses allow embryos of annual killifish to survive dry periods that may occasionally span even multiple years. Egg provisioning thus represents a crucial life-history process in annual killifish that may affect embryonic survival or amount of energetic reserves available after hatching.

Bigger annual killifish species indeed produce disproportionately larger eggs compared to same-sized non-annual species (Eckerström-Liedholm et al., 2017). In the African annual killifish *Nothobranchius furzeri* (Nothobranchiidae), egg size determines hatchling size irrespective of the time spent in incubation (Vrtílek et al., 2017). This suggests that some resources in the egg that scale with egg size affect hatchling size and that these are not consumed during diapause. There is not much data from other annual killifish species, but Moshgani and Van Dooren (2011) found substantial within-species variation in egg size of the South-American annual *Austrolebias nigripinnis* (Rivulidae). They did not, however, relate egg size to amounts of resources or to hatchling size.

Here, we test for differences in allometries of egg content parameters with egg size in ecologically and phylogenetically divergent species of annual killifish from South America (genus *Austrolebias*) and Africa (species *N. furzeri*). We combine analyses of different egg composition parameters – wet and dry egg mass; carbon, nitrogen and sulphur content; the amount of triglycerides – to quantify resources such as water, nutrients and energy in individual eggs. We aimed to test the relevance of egg area measured from digital photographs as a predictor for egg wet and dry weight within and across annual killifish species. In addition to that, we incubated a subset of the collected clutches to assess the effect of egg area on hatchling traits. We evaluate the generality of our conclusions by using two additional independently collected datasets.

## Methods

### Breeding pairs of *Austrolebias* species

To collect eggs, we established breeding pairs of four *Austrolebias* species from captive populations maintained at the CEREEP station in Nemours-St. Pierre, France (approval no. B77-431-1). We used two small species – *A. bellottii* (population “Ingeniero Maschwitz”, 4 pairs) and *A. nigripinnis* (“La Guarderia”, 4 pairs); and two large (“piscivorous”) species – *A. elongatus* (“General Conesa”, 5 pairs) and *A. prognathus* (“Salamanca”, 2 pairs). Table S1 in the Supplementary Information lists the ages and body sizes of the parental fish and includes details on their source populations.

**Table 1.** Overview of species mean values for the measured egg parameters. The “Females” column reports number of individual spawning females whose eggs were analysed for composition or went into incubation (the number in brackets corresponds to females in the incubation dataset if the two differ). The numbers for egg parameters shown in brackets give sample sizes. Standard deviation (SD) is provided for egg area along with the mean.

| Species | Females | Egg area ± SD [mm<sup>2</sup>] | Water<br>content [%] | N\|C\|S content [%] | Triglyceride<br>concentration [mg/ml] | Incubated/<br>Hatched |
| --- | --- | --- | --- | --- | --- | --- |
| *Austrolebias bellottii* (small species) (AB) | 4 | 1.336 ± 0.188 (928) | 68.5 (99) | 8.5\|54.4\|0.4 (39) | 6.8 (10) | 829/39 |
| *Austrolebias nigripinnis* (small species) (AN) | 4 | 1.214 ± 0.148 (478) | 68.5 (63) | 8.6\|54.9\|0.4 (28) | 4.7 (3) | 415/51 |
| *Austrolebias elongatus* (piscivore) (AE) | 2 (5) | 3.781 ± 0.515 (116) | 74.8 (14) | 9.1\|51.9\|0.5 (12) | 16.8 (6) | 102/2 |
| *Austrolebias prognathus* (piscivore) (AP) | 1 (2) | 4.420 ± 0.589 (49) | 73.4 (11) | 8.8\|52.4\|0.5 (11) | 12.9 (4) | 38/0 |
| *Nothobranchius furzeri* (small species - African) (NF) | 7 | 1.154 ± 0.105 (108) | 73.5 (108) | 8.9\|54.3\|0.4 (40) | 3.9 (10) | - |
| *Austrolebias bellottii* 6+ years in incubation (AB 2011) | - | 1.689 ± 0.176 (7) | - | - | 9.9 (7) | - |
| Additional datasets |  |  |  |  |  |  |
| *Austrolebias bellottii* 2015 (AB 2015) | 3 | 1.357 ± 0.370 (61) | 83.1 (61) | - | - | - |
| *Austrolebias nigripinnis* 2015 (AN 2015) | 1 | 1.594 ± 0.118 (28) | - | - | - | -/28 |
| *Austrolebias elongatus* 2015 (AE 2015) | 1 | 3.813 ± 0.524 (23) | - | - | - | -/23 |

The breeding pairs were kept in tanks of 21 litres for the small species or of 54 l for the piscivores in a dedicated climate room at the CEREEP facility. To prepare tank water, we mixed reverse osmosis and conditioned (JBL NitratEx, PhosEx, SilicatEx resin-filtered and Sera Toxivec 0.5 ml/l) tap water in a 1:1 ratio resulting in 200-300 µS conductivity. We replaced 1/3 of the water per tank on a weekly basis. The room temperature was set to 23 °C during the light period of the day (8AM-10PM, 14 h) and to 18 °C when dark (10PM-8AM, 10 h), making water temperature vary between 17-23 °C. Fish were fed daily with live *Tubifex* sp. (Grebil Pére et Fils, Arry, France).

### Breeding stock of *N. furzeri*

We wanted to compare egg content between two phylogenetically distinct genera of annual killifish - *Austrolebias* (South America) and *Nothobranchius* (Africa) (Furness et al., 2015). We therefore established a breeding stock of *N. furzeri* (population “MZCS414”) from the Institute of Vertebrate Biology CAS in Brno, Czech Republic at the CEREEP facility. We hatched *N. furzeri* according to the standard protocol (Polačik et al., 2016) using a 1:1 mixture of osmotic and tap water at 18 °C. We subsequently reared the fish at 27-28 °C and fed them with *Artemia salina* nauplii until the age of ten days when they were weaned on *Tubifex* sp. We separated sexes when first marks of nuptial colouration of males appeared, at approximately three weeks, and paired the fish only for egg collection. *Nothobranchius furzeri* were hatched on July 4, 2017, shortly after the collection of eggs from *Austrolebias* began, so they reached full maturity (at an age of six weeks) when *Austrolebias* egg collection finished. The breeding stock of *N. furzeri* consisted of seven pairs.

### Egg collection and measurements

We collected *Austrolebias* eggs using two- or five-litre plastic spawning containers (depending on the species). The containers were filled with a 0.5 cm layer of 300-400 µm diameter glass beads and 5 cm of peat granules (2-4-mm diameter fraction; HS aqua Torogran, Ulestraten, Netherlands) to allow fish to dive into the substrate during spawning. Each tank had a single spawning container which we replaced every other evening. Retrieved containers were left overnight in an adjacent climate room at 23 °C. We collected eggs the following day using set of sieves of different mesh sizes as described in (Moshgani and Van Dooren, 2011). We paired *Nothobranchius* to spawn for two hours (9-11AM) in two-litre containers with one cm layer of the glass beads. We collected eggs six hours later by sieving the spawning substrate.

We photographed fertilized killifish eggs and only measured eggs with a clear perivitelline space. Eggs were gently cleaned with a paintbrush and we photographed (Nikon COOLPIX 4500) ten random eggs per pair at each collection date under a dissecting microscope (Olympus SZH10) at 30x magnification. We measured two egg dimensions: the longest egg axis (*d*1) and the perpendicular axis (*d*2) in pixels (ImageJ ver.1.50i) and subsequently rescaled to lengths in millimetre using a 0.01-mm calibration slide. From these two lengths, *egg area* was calculated as 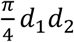. We use mainly egg area as the measure of egg size in this study.

### Preparations for the analysis of egg content

We collected freshly laid eggs from *Austrolebias* species for the analysis of egg content at four occasions among the egg collections for incubation (see “Incubation of embryos and hatching” below). After photographing, we stored each egg in a separate 1-ml Eppendorf tube with UV-sterilized water (mixed as for incubation) at 23 °C in the incubator. Next day, we removed moisture from the eggs by rolling them over a filter paper and measured their wet weight to the nearest 0.001 mg (Mettler Toledo XPE26 balance). The eggs for analysis of carbon, nitrogen and sulphur content (see “Elemental content analysis” below) were subsequently oven-dried for 24 h at 70 °C, weighed for dry weight and stored at 4 °C. The eggs for the triglyceride assay (see “Triglyceride assay” below) were each put fresh in a dry Eppendorf tube and frozen at -20 °C before the analysis.

We took the opportunity to get a sample of eggs from extremely old clutches of *A. bellottii* spawned in February 2011 (6+ years of incubation) and compared their triglyceride content to that of freshly laid eggs of the same species. These clutches had been stored in damp mixed coco-coir and peat in zip-locked bags. We briefly dried the substrate to make the eggs visible, put the retrieved eggs into UV-sterilized water and then treated them in the same way as those above for the triglyceride assay.

### Elemental content analysis

The concentrations (weight percentage) of carbon, nitrogen and sulphur were determined by dry combustion with a CNS element analyser (Thermo Scientific Flash 2000) at the Global Change Research Institute CAS, Brno, Czech Republic. Benzoic acid was used as a reference material.

### Triglyceride assay

We assessed egg triglyceride reserves using a commercial Triglyceride Colorimetry Assay kit (ref. 10010303; Cayman Chemical, Ann Arbor, MI, USA) following the protocol for tissue homogenates. We left frozen eggs to thaw at room temperature and put each one in a 1.5-ml Eppendorf tube with 0.2 ml of 5X salt solution (Standard Diluent). The tube content was homogenized using a disperser (IKA T10 basic ULTRA-TURRAX) for 30 s and then centrifuged for 10 min (Eppendorf 5430, 10,000 r/min, 4 °C) to separate egg envelope and content. We put the supernatant into a clean Eppendorf tube and mixed briefly on a vortex (Fisherbrand L-46). We pipetted 10 µl aliquot in a prepared 96-well plate, added 150 µl of diluted Enzyme Mixture and incubated the plate at room temperature for 10 min. We made two replicates per sample. We measured the absorbance at 540 nm wavelength three times per replicate using a microplate reader (BIO-RAD iMark). We used the reference standards to calculate triglyceride concentrations of the two replicates.

### Incubation of embryos and hatching

To test for long-term effects of egg size on embryonic development, we incubated a sample of the photographed eggs and hatched those having completed their development after a pre-determined duration. We stored eggs individually in 24-well plates filled with UV-sterilized mixed water (1:1 ratio of aged tap and reverse osmosis water) at 23 °C in a Memmert IPP500 incubator. We inspected development and survival of incubated embryos weekly. Inspections were more frequent, however, during the first two weeks when embryonic mortality was highest. At each inspection, we removed dead embryos and recorded the date of those reaching the pre-hatching developmental stage, i.e. the “golden-eye” stage (Stage 5 in Varela-Lasheras and Van Dooren (2014), St. 43 in Podrabsky et al. (2017)). We kept the pre-hatching stage embryos in their wells until the scheduled hatching date. A minor fraction of the pre-hatching stage embryos (7 %, 29 of 427), however, hatched spontaneously prior to the planned hatching date. These individuals were not used in further analyses of hatchling size.

We scheduled hatching four months after the date of last spawning and maximum time spent in the pre-hatching stage was 92 days. To initiate hatching, we gently wiped egg dry by rolling over a filter paper and put it back in a dry well. We stimulated hatching by rewetting the embryos 3 h later using a 6:1 mixture of 23 °C osmotic water and peat extract (1 l of peat boiled in 1 l of osmotic water). We inspected hatching success eight hours after rewetting, anaesthetized the hatched fish with clove oil and photographed them from a lateral view under a dissecting microscope (magnification 15x for small species and 10x for piscivores). We measured standard length (SL) of each hatchling and the area of its remaining yolk. We repeated the hatching procedure on the following day with the previously unhatched embryos.

### Independent datasets of *Austrolebias* egg size, wet and egg dry weight, and hatchling size

We used two independent datasets of *Austrolebias* eggs collected in 2015 to generalize our findings. The first dataset consisted of eggs of *Austrolebias bellottii* (population “Ingeniero Maschwitz”, 3 pairs, 61 eggs), where egg area was measured together with wet and egg dry weight. The eggs were collected from spawning boxes containing moss *Sphagnum magellanicum* facilitating rapid egg retrieval. We measured egg area, egg wet and dry weight as described above.

The other independent dataset was on egg size-hatchling size relationship from a pair of *A. elongatus* (“Vivorata”) and of *A. nigripinnis* (“San Javier Missiones”) kept in a greenhouse. We collected eggs from spawning boxes with a mix of coco-coir and peat during February and March 2015. The substrate was gently dried until eggs could be seen and collected. Again, we measured egg size as above and transferred eggs to 24-well plates with UV-sterilized water at two different temperatures (22 °C and 25 °C) for incubation. Three months later (May 28, 2015), embryos in the pre-hatching stage were air dried for four hours and rewetted to initiate hatching, so we could measure standard length (SL) of the hatchlings.

### Statistical analysis

We used the data for two different but related purposes: (1) we determined which explanatory variables contributed significantly to variability in egg parameters with a particular focus on species differences in allometric scaling exponents of egg size; and (2) we assessed how well egg size, measured as egg area on digital images, predicted egg wet weight, dry weight and hatchling size.

We assumed that the investigated egg parameters scaled allometrically with egg size and that these relationships can be described by constants of proportionality and scaling exponents (Appendix, Section A). We used egg area as a measure of size. In the Appendix, we explain how allometries can be analysed in a multi-species context and provide arguments for the modelling choices we made. We ignore effects of measurement error in egg size in our analysis (Appendix, Section F).

The parameters we measured in eggs are expected to covary (water weight with egg dry weight, for example) both between and within species and we therefore used multivariate models wherever possible (Appendix, Section D). To do so, we reformatted values of response variables into stacked form and added a dummy factor “trait” with levels identifying the different egg parameters. All models thus contained “trait” fixed effects by default to distinguish the egg parameters and to allow for fitting trait-specific random effects and different residual variances per trait. We fitted models using functions from library “nlme” (Pinheiro and Bates, 2000) in R software v. 3.5.3 (R Core Team, 2019).

We always started model selection with a full model and then tried to simplify it on the basis of log-likelihood ratio tests and F-tests on model log-likelihoods. We removed random effects first, followed by simplification of residual variance structures and then fixed effects (Zuur et al., 2009). We removed random effects by comparing models containing all residual variance terms and fixed effects fitted with restricted maximum likelihood (REML). When testing the importance of individual fixed effects, we refitted the models with maximum likelihood (ML). We then assessed fit of the selected models by inspecting the distributions of residuals.

To test species-specific allometries in egg dry mass and water content related to egg size, we used egg dry weight and water weight as log-transformed response variables in a bivariate model. Egg water weight was fitted as egg wet weight minus dry weight, therefore we did not add total wet weight as an explanatory variable (Appendix, Section E). The full model included “trait”, “species” and “log-transformed egg area” with all the possible interactions as fixed effects to account for different allometries between species and traits. The log-transformed egg area was zero-centered per species prior the analysis to ensure that intercept difference terms capture the total differences between groups while slopes estimate the scaling exponents (Appendix, Section D; Nakagawa et al., 2017). We provide a simple rule here which can help interpreting estimated scaling exponents. If egg parameters scale isometrically with egg volume and eggs are spheres, then they must scale in a hyperallometric manner with egg area (with coefficient 3/2, Appendix, Section B). Hence, when exponents of egg area are not estimated to be above 3/2, there are no hyperallometries for volume. Parental pair and collection date were modelled as trait-specific random effects and residual variances were allowed to differ between species and traits.

Carbon, nitrogen, and sulphur (CNS) are components of egg dry weight with raw data values given as percentages. We carried out compositional analysis with CNS percentages transformed using isometric log-ratio (ilr) (library “compositions”; van den Boogaart and Tolosana-Delgado, 2013). We fitted them as responses in a trivariate general linear model because mixed-effects model did not converge. The full model was of the same structure as the bivariate model above for egg dry weight and water weight. Individual model parameters in the compositional analysis were, however, difficult to interpret (van den Boogaart and Tolosana-Delgado, 2013). We therefore report parameter estimates from the best model fitted with log-transformed total amounts of individual elements calculated from CNS percentages and dry weights (Appendix, Section E).

We assayed triglyceride content to test for relative species differences. Triglyceride content was measured as diluted sample concentration in two replicates from the same egg. We then back-calculated the total amount of egg triglycerides in the replicates using the sum of standard solvent volume and egg volume taken as 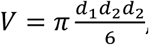, assuming a spheroid egg shape. We used log transformed total amount of triglycerides from each replicate as a response and corrected for different egg sizes by fitting log-transformed egg volume as an offset (Appendix, Section E). We fitted fixed effect of “species” to test for intercept differences among species and interaction between “species” and species-centered “log-transformed egg volume”. Full mixed-effects model that would control for both pseudoreplication and species-specific residual variances did not converge. We therefore fitted two alternative models: a mixed-effects model with sample ID as random effect; or a general linear model with averaged values from the two subsamples accommodating species-specific residual variance. The outcome of these two models was qualitatively similar and we report estimates of species contrasts from the model with the averaged sub-sample values based on Tukey’s post-hoc test (function glht(), library “multcomp”; Hothorn et al., 2008).

We tested for long-lasting effects of egg size on traits at hatching: hatchling size and remaining yolk reserves. Hatchling size and yolk sac area were log-transformed and analysed using a bivariate mixed-effects model as above for egg dry weight and water content. The full model included a triple interaction between “trait” (by default), “species” and species-centered “log-transformed egg area” to account for different allometries in species and traits (Appendix, Section C). In addition to that, we fitted an interaction between “trait” and the number of days spent in the pre-hatching stage. This was to account for the variation in pre-hatching stage duration among embryos. Full mixed-effects model containing random effects of pair and collection date did not converge. We thus fitted general linear models with trait-specific residual variances.

To evaluate how well egg area predicts egg wet weight, dry weight, and hatchling size, we calculated coefficients of determination (*R^2^*) in mixed-effects models using function r.squaredGLMM() (library “MuMIn”; Bartoń, 2018). This function allows to separate *R^2^* for the whole model 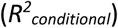 from the fixed-effects part of the model 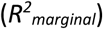 (Nakagawa and Schielzeth, 2013). We first fitted full mixed-effects model with “species” in interaction with untransformed “egg area” as fixed effects (Appendix, Section G), and pair and collection date random effects. Residual variances were allowed to vary between species. We then fitted separate models per species with egg area fixed effects, and pair and collection date random effects. We fitted the same models to the additional datasets from 2015 to compare predictive power of egg area among different studies.

## Results

We collected over six thousand eggs of five annual killifish species in 2017 of which we incubated 1384 and measured wet and dry weight of 295 eggs. We used 130 of these dried eggs for CNS elemental analysis and another 40 fresh eggs, which were weighed, for triglyceride content assay (including seven eggs collected from substrate incubated for 6+ years) (Table 1). The additional two datasets consisted of 61 wet and egg dry weights from a single species, and of 51 hatchling sizes from another two species measured in 2015 (see Table 1 for more details).

### Egg size

Taking egg area determined from images as the measure of egg size, the collected eggs clustered into two groups according to adult sizes of the species they belonged to. Eggs were notably larger in the piscivores, *Austrolebias elongatus* and *A. prognathus*, compared to the small *Austrolebias* species, *A. bellottii* and *A. nigripinnis*, and to *Nothobranchius furzeri* (Fig. 1, Table 1). The size of *A. bellottii* eggs collected after 6+ years of incubation was larger compared to freshly laid eggs of the same species (Table 1). The eggs from the additional datasets were similar in size to eggs collected for this study (Fig. 1).

**Fig. 1.**
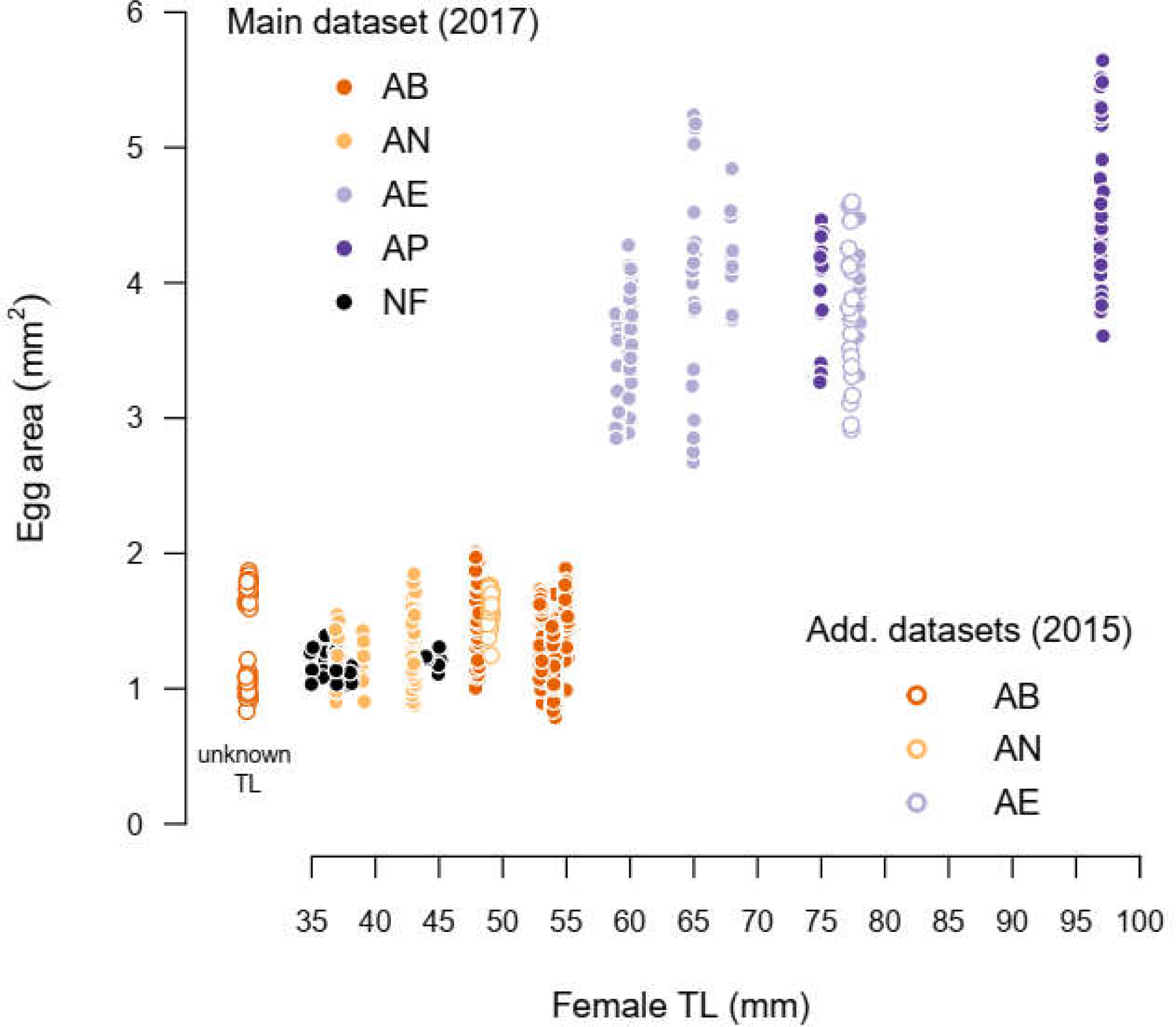
Species with larger body size lay larger eggs. The plot shows the relationship between female body size and egg size in the studied species of annual killifish. Full points represent values collected for this study and empty points are for data from the two additional datasets. Note that for data obtained from *Austrolebias bellottii* in 2015 size of the three females was unknown. Species abbreviations: AB – *Austrolebias bellottii*, AN – *A. nigripinnis*, AE – *A. elongatus*, AP – *A. prognathus*, NF – *Nothobranchius furzeri*.

### Dry egg mass and water content

The allometric scaling exponents of egg dry and water weight on egg area differed between species. The best model included a three-way interaction between species, trait and log-transformed egg area (species:trait:log(egg area) interaction, *F*4,532 = 2.67, *p* = 0.032). This means that the effects of log-egg area on log-egg dry weight and log-water weight differed and that this difference depends on the species (Table 2, Fig. 2A). However, the water weight allometric scaling exponent was significantly different from *A. bellottii* and from dry weight only in eggs of *A. prognathus.* In *A. prognathus,* water weight appeared to scale isometrically with egg area while egg area had no effect on egg dry weight (Fig. 2A). *Austrolebias bellottii* and *A. nigripinnis* showed hypo-allometric, and *A. elongatus* hyper-allometric effects of egg area on dry weight. None of the predicted exponents were above 3/2 (Table 2), hence there were no hyperallometries for the relationship with egg volume. The estimates for dry weight effects in *A. prognathus* and *N. furzeri* were close to zero indicating very weak relationships between egg size and dry weight. In the additional 2015 dataset, *A. bellottii* eggs’ dry weight changed hyperallometrically with egg area (log(egg area) effect ± SE (standard error): 1.360 ± 0.085). Water weight changed similarly to that (difference between log(egg area) effect on log(dry weight) vs. log(water weight): -0.239 ± 0.128), hence both changed approximately isometrically with egg volume.

**Fig. 2.**
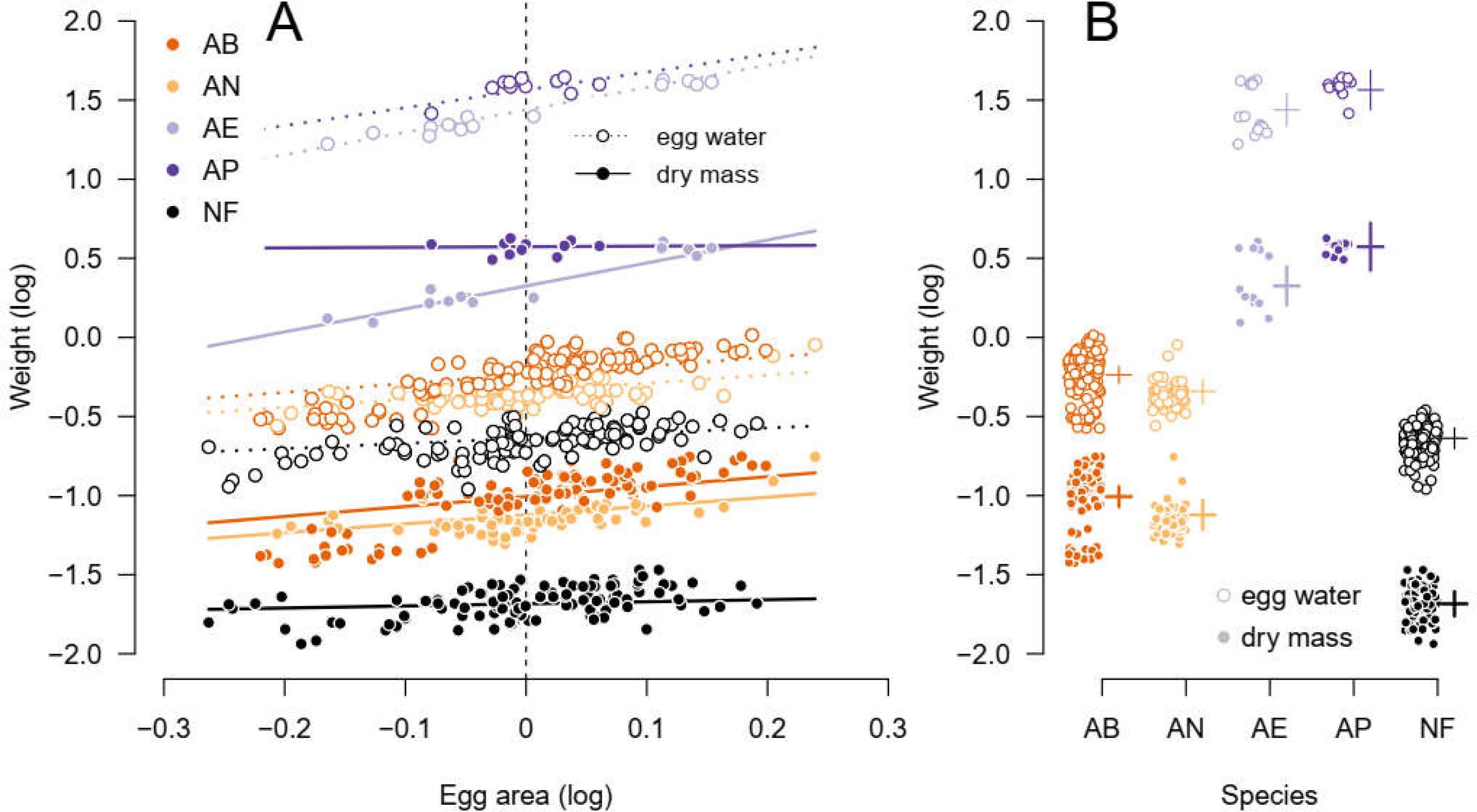
Species-specific allometries between egg size and egg dry mass, and egg size and water content. Plot (A) shows species-specific slopes of the relationship between log-dry weight and the species zero-centered log-egg area (full points, solid lines), and log-water weight and species zero-centered log-egg area (empty points, dashed lines) (Table 2). Note that the lines for egg dry weight and water weight are parallel in most of the species except *A. prognathus* (AP). The parallel lines mean that water content scaled similarly to dry mass with egg size. Plot (B) then shows relative amount of water and dry mass in the eggs of the different species. The distance between the log-dry and log-water weight here corresponds to log-ratio of these two egg parameters (means are denoted by horizontal lines and vertical lines give standard errors estimated from the model, Table 2). For their mean egg size (indicated as vertical dashed line at 0 in Figure 2A), the piscivores (*A. elongatus* and *A. prognathus*) and *N. furzeri* have relatively higher proportion of water weight to dry mass (wider distance between mean log-dry weight and mean log-water weight). Species abbreviations: AB – *Austrolebias bellottii*, AN – *A. nigripinnis*, AE – *A. elongatus*, AP – *A. prognathus*, NF – *Nothobranchius furzeri*.

**Table 2.** Species-specific allometries of egg dry mass and water content with egg area. We zero-centered the log-transformed egg area for each species prior to analysis. The “log(dry weight)” column corresponds to the mean effects of species on log-transformed egg dry weight, where a first set of species effects represents differences from *A. bellottii*. The species differences for log-egg area effects (lower half of the first column) are differences in slopes of log-dry weight on centered log-egg area from that of *A. bellottii* (Fig. 2A). The “log(water weight) vs log(dry weight)” column gives estimates of the mean multiplicative species differences in log-water weight compared to *A. bellottii.* The reference estimate for *A. bellottii* here represents the log-ratio between egg water weight and dry weight (Fig. 2B). Note that egg water weight approximately increased with egg area (lower half of the second column) across four out of five species, with the same scaling exponent as egg dry weight. Parameter estimates are given with their standard errors (SE). Estimates significantly different from zero (*p* < 0.05) are in bold.

| Fixed Effect | log(dry weight) | log(water weight) vs log(dry weight) |
| --- | --- | --- |
| *A. bellottii* (reference) | -1.006 (0.033) | 0.769 (0.021) |
| Difference of *A. nigripinnis* | **-0.116** (0.048) | 0.012 (0.032) |
| Difference of *A. elongatus* | **1.332** (0.063) | **0.343** (0.043) |
| Difference of *A. prognathus* | **1.579** (0.077) | **0.222** (0.051) |
| Difference of *N. furzeri* | **-0.678** (0.040) | **0.277** (0.028) |
| log(egg area) (effect in *A. bellottii*) | **0.628** (0.116) | -0.074 (0.134) |
| Difference of log(egg area) in *A. nigripinnis* | -0.066 (0.155) | 0.022 (0.183) |
| Difference of log(egg area) in *A. elongatus* | **0.823** (0.180) | 0.038 (0.216) |
| Difference of log(egg area) in *A. prognathus* | -0.592 (0.321) | **1.164** (0.401) |
| Difference of log(egg area) in *N. furzeri* | **-0.496** (0.156) | 0.268 (0.183) |

Overall, the annual killifish eggs contained between two and three times as much water as dry matter. The estimated differences in species intercepts or of different traits on the same species represent multiplicative total differences after back-transformation (Fig. 2 Table 1; Appendix, Section D). Relative content of water in the eggs of *A. elongatus, A. prognathus*, together with *N. furzeri* was higher than in the small *Austrolebias* species, *A. bellottii* and *A. nigripinnis* (Tables 1 and 2, Fig. 2B). Eggs of *A. bellottii* from 2015 contained 2.66-times more water than dry matter (log-transformed estimate: 0.979 ± 0.035), compared to 2.16 ratio in *A. bellottii* eggs collected in 2017. Egg dry weight appeared to vary more than water weight (estimated SDs ratio between log(dry weight) and log(water weight) was 1.294; CI [1.140, 1.467]) and eggs of *N. furzeri* were the most variable (SD (standard deviation) ratio was 1.723; CI (confidence interval) [1.256, 2.362]).

### CNS elemental analysis

Carbon, nitrogen and sulphur contents varied among eggs of different species including species-specific allometric scaling exponents of egg area (Table 3). The compositional analysis with ilr-transformed data showed a significant three-way interaction (species:trait:log(egg area), *F8,360* = 2.20, *p* = 0.028) meaning that proportions of the three elements scaled differently with egg area and that these differences depended on the species. As the estimates from ilr-transformed data were hard to interpret, we re-fitted the full model with log-transformed total amounts of the elements (Table 3). Overall, carbon content scaling exponents of egg area varied among species, but nitrogen and sulphur scaling exponents were hardly different. The only exception was the scaling exponent estimate for sulphur in eggs of *A. nigripinnis* (Table 3). None of the slopes of egg area is significantly above 1.5, hence we can again conclude that there are no hyperallometries for egg volume.

**Table 3.**
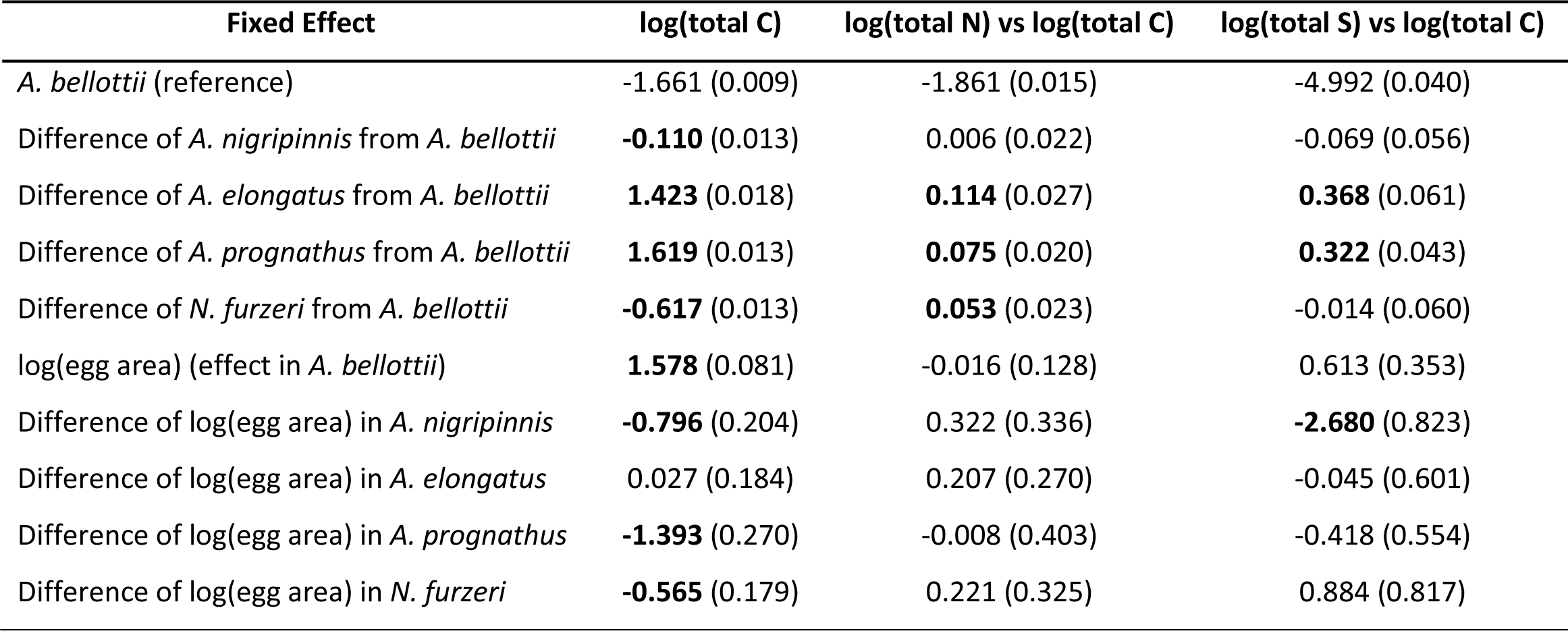
Relative changes in egg total amounts of carbon, nitrogen and sulphur between species. The “log(total C)” column gives the estimate of mean log-transformed total egg carbon for each species using *A. bellottii* as a reference followed by slopes of centered log-egg area. The two other columns correspond to species differences between log-total nitrogen and log-total carbon, and log-total sulphur and log-total carbon, respectively. Relative elemental composition of eggs varied across different species. The top half of the table indicates that the piscivores’ (*A. elongatus* and *A. prognathus*) and *N. furzeri* eggs had relatively more nitrogen to carbon than *A. bellottii* (Fig. 3A). In addition to that, eggs of piscivores also contained relatively more sulphur (to carbon) (Fig. 3B). Allometric scaling of elements content with egg area differed among species, but was consistent for the different elements (except for sulphur content in *A. nigripinnis*). The estimates are given with their standard errors (SE). Values significantly different from zero (*p* < 0.05) are in bold.

| Fixed Effect | log(total C) | log(total N) vs log(total C) | log(total S) vs log(total C) |
| --- | --- | --- | --- |
| *A. bellottii* (reference) | -1.661 (0.009) | -1.861 (0.015) | -4.992 (0.040) |
| Difference of *A. nigripinnis* from *A. bellottii* | **-0.110** (0.013) | 0.006 (0.022) | -0.069 (0.056) |
| Difference of *A. elongatus* from *A. bellottii* | **1.423** (0.018) | **0.114** (0.027) | **0.368** (0.061) |
| Difference of *A. prognathus* from *A. bellottii* | **1.619** (0.013) | **0.075** (0.020) | **0.322** (0.043) |
| Difference of *N. furzeri* from *A. bellottii* | **-0.617** (0.013) | **0.053** (0.023) | -0.014 (0.060) |
| log(egg area) (effect in *A. bellottii*) | **1.578** (0.081) | -0.016 (0.128) | 0.613 (0.353) |
| Difference of log(egg area) in *A. nigripinnis* | **-0.796** (0.204) | 0.322 (0.336) | **-2.680** (0.823) |
| Difference of log(egg area) in *A. elongatus* | 0.027 (0.184) | 0.207 (0.270) | -0.045 (0.601) |
| Difference of log(egg area) in *A. prognathus* | **-1.393** (0.270) | -0.008 (0.403) | -0.418 (0.554) |
| Difference of log(egg area) in *N. furzeri* | **-0.565** (0.179) | 0.221 (0.325) | 0.884 (0.817) |

Egg elemental composition varied across different species. Species differences in the total amount of carbon can be largely explained by species effects on egg dry weight (compare the “log(dry weight)” column of Table 2 to the “log(total C)” in Table 3). The top half of the table indicates that the piscivores (*A. elongatus* and *A. prognathus*) and *N. furzeri* had relatively more nitrogen to carbon than *bellottii* (Fig. 3A). In addition to that, eggs of piscivores also contained relatively more sulphur (to carbon) (Fig. 3B, Table 3). The three elements analysed together made up between 60 and 68 % of egg dry weight (carbon 54.1 %, nitrogen 8.7 % and sulphur 0.4 %, on average; Table 1). The variance was particularly high for measurements of sulphur in eggs of the small species (*A. bellottii*, *A. nigripinnis* and *N. furzeri*) (SD ratios were 6.330-8.980).

**Fig. 3.**
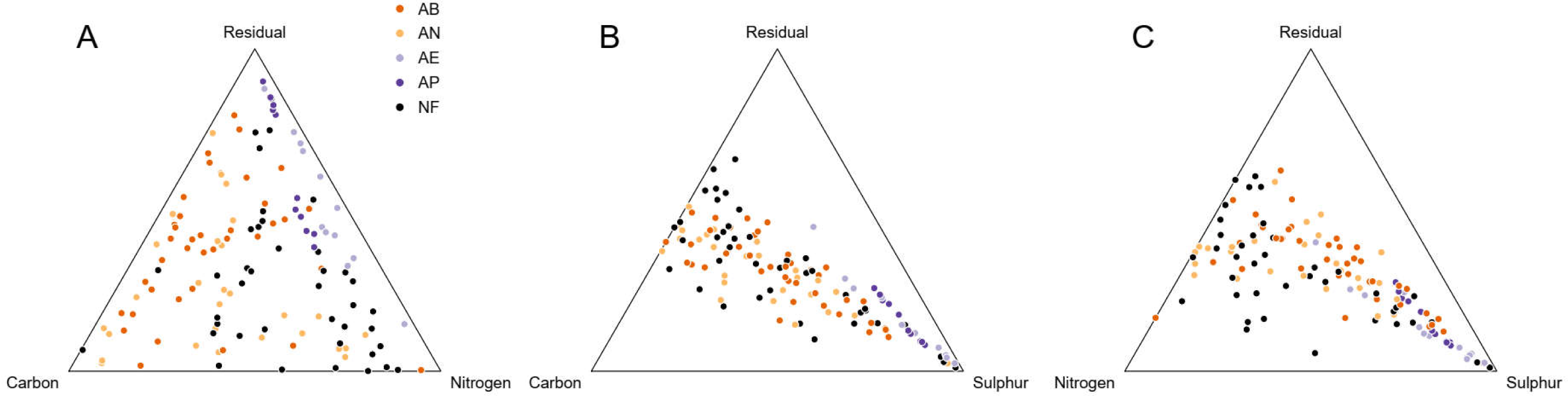
Egg elemental composition differed among species. The ternary plots show relationship between the three measured elements in killifish eggs and the residual content (A – carbon, nitrogen and residual; B – carbon, sulphur and residual; C – nitrogen, sulphur and residual). The position of each point (trivariate coordinate for each egg) reflects the inter-dependence of the elemental content proportions as they together must add up to 100 %. The original percentage values were transformed using isometric log-ratio transformation, and then scaled and centered for visualization. Species abbreviations: AB – *Austrolebias bellottii*, AN – *A. nigripinnis*, AE – *A. elongatus*, AP – *A. prognathus*, NF – *Nothobranchius furzeri*.

### Triglyceride content analysis

The relative amount of egg triglycerides varied within and among species (species, *F5,34* = 12.88, *p* < 0.001; Fig. 4). We found that, after accounting for different egg volumes, the amount of egg triglycerides was highest in *A. elongatus* and lowest in *N. furzeri* which overlapped with the small *Austrolebias* species, *A. bellottii* and *A. nigripinnis* (Fig. 4). We didn’t observe a difference between fresh eggs of *A. bellottii* and those collected from 6+ years old substrate (Fig. 4). We explored potential allometric scaling exponent effects of egg volume on the amount of triglycerides and found a significant species (species:log(egg volume) interaction, *F6,28* = 3.35, *p* = 0.013) where *N. furzeri* had a much larger positive value than the other species. However, this significant interaction and the large estimate are due to four very small outlying concentrations among the smallest eggs, suggesting that these eggs were incompletely composed and that we should not take this as evidence for hyperallometry.

**Fig. 4.**
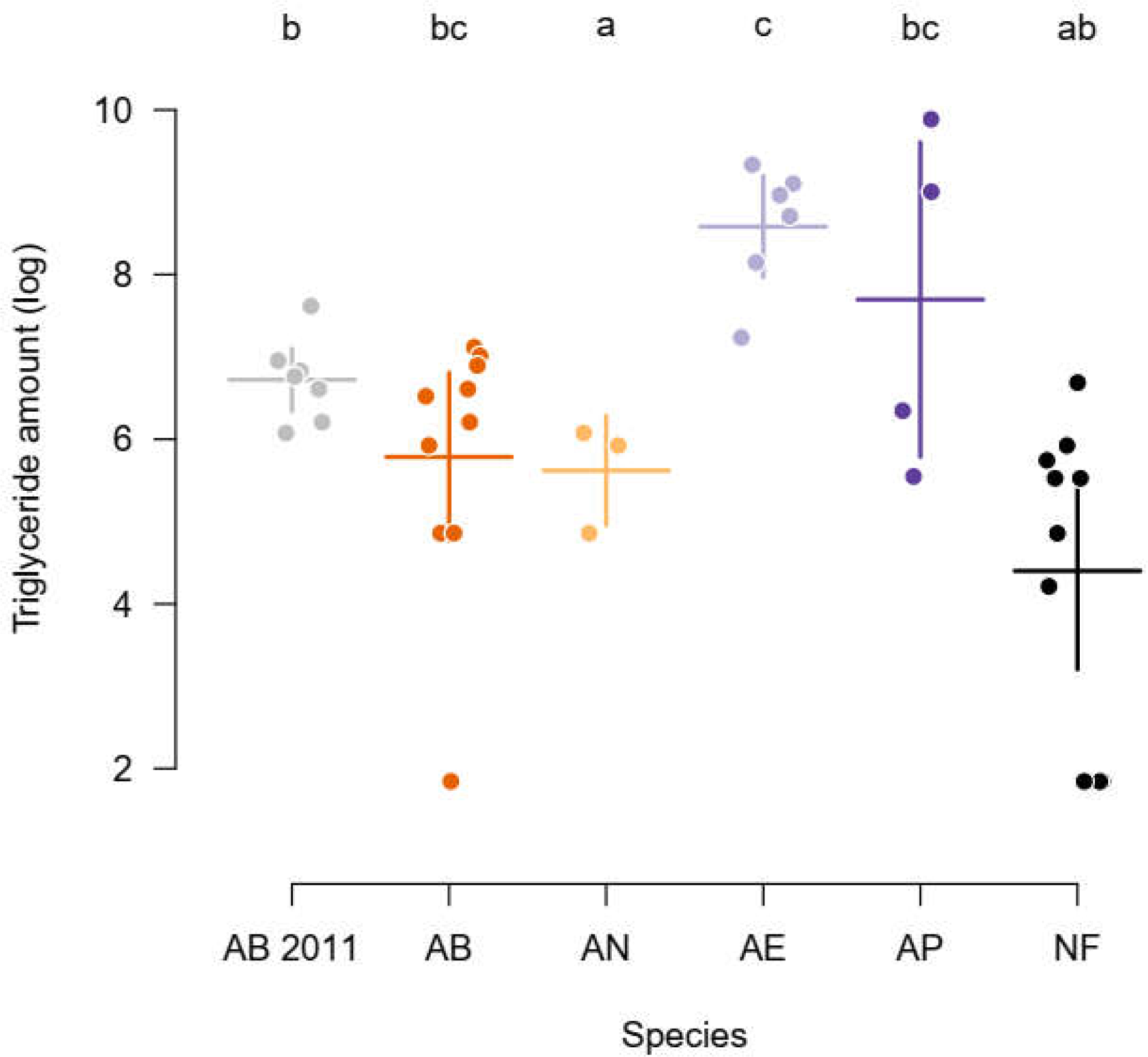
Relative triglyceride content was higher in species with larger eggs. Note that the extremely long incubation (6+ years) did not reduce the level of triglycerides in the eggs of *Austrolebias bellottii* from 2011 compared to the freshly laid eggs of the same species. Horizontal lines are for species mean and vertical lines for standard errors from the model. The lowercase letters on top of each species data indicate differences in the relative amount of triglycerides (based on a model that included species zero-centered log-egg volume as an offset). Each point represents log-transformed amount of triglyceride amount in an egg. Species abbreviations: AB – *Austrolebias bellottii*, AN – *A. nigripinnis*, AE – *A. elongatus*, AP – *A. prognathus*, NF – *Nothobranchius furzeri*.

### Hatchling size and remaining yolk

We successfully hatched 22% (92/427) of pre-hatching stage embryos. *Austrolebias bellottii* hatchlings were larger than *A. nigripinnis* (Table 4). Within species, hatchling size did not depend on egg area, but yolk sac area decreased with egg area (trait: centered log(area) interaction, *F1,172* = 4.73, *p* = 0.031, Fig. 5, Table 4). Yolk sac area also diminished with the number of days embryos spent in the pre-hatching stage and we did not record any gain to the embryo by growing larger while waiting for hatching (trait:daysPH interaction, *F1,172* = 4.61, *p* = 0.033). In the additional 2015 dataset, there was again an at most weak effect of egg area on hatchling size of *A. nigripinnis* (log(egg area) effect: 0.165 ± 0.091), hatchlings of *A. elongatus* showed a hypo-allometric increase in size with egg area (log(egg area) effect: 0.381 ± 0.057) (Fig. 5). Yolk sac size had a higher variance than hatchling size in our 2017 dataset (estimated yolk sac size:hatchling size SDs ratio was 5.291; CI [4.304, 6.506]).

**Fig. 5.**
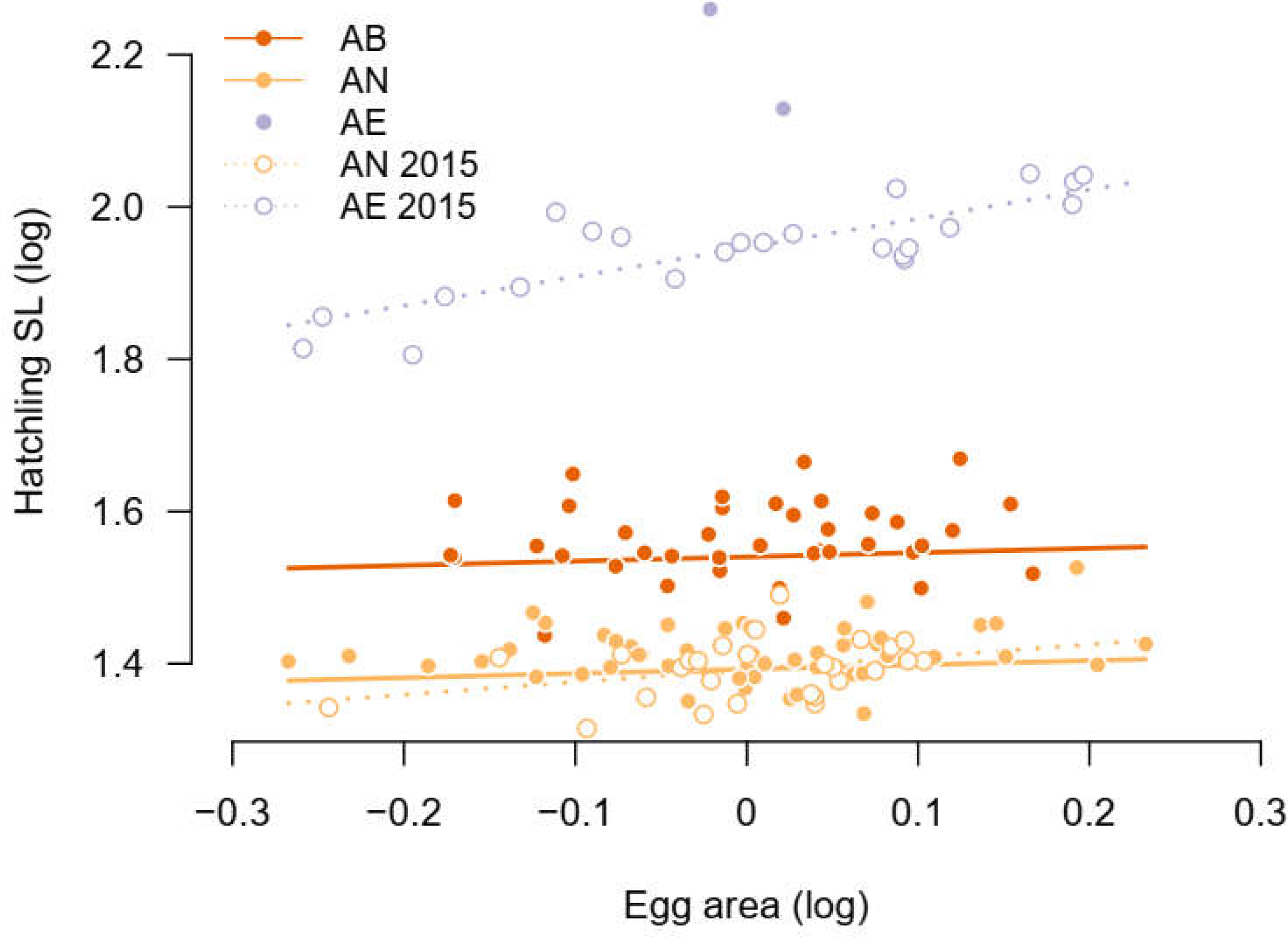
Hatchling size corresponds with egg size only in the large species. Hatchlings coming from larger eggs were bigger only in *A. elongatus* (piscivore) from 2015, but in the small *Austrolebias* (*A. bellottii* and *A. nigripinnis*) egg area had no effect on hatchling size. Species abbreviations: AB – *Austrolebias bellottii*, AN – *A. nigripinnis*, AE – *A. elongatus*.

**Table 4.** Effect of egg size and duration of pre-hatching stage on hatchling size and yolk sac size. The “log(hatchling size)” column reports parameter estimates of species differences and scaling exponents of egg area (Fig. 5). The “log(yolk sac area) vs log(hatchling size)” column then gives the proportional changes for yolk sac area. The effect of log-transformed egg area appeared to be zero on hatchling size, but negative on yolk sac area in both of the species. Yolk sac area declined with the number of days spent in the pre-hatching stage. Estimates are given with their standard errors (SE). Terms significantly different from zero (*p* < 0.05) are in bold.

| Fixed Effect | log(hatchling size) | log(yolk sac area) vs log(hatchling size) |
| --- | --- | --- |
| *A. bellottii* (reference) | 1.540 (0.025) | 3.437 (0.136) |
| Difference of *A. nigripinnis* from *A. bellottii* | **-0.148 (0.009)** | -0.091 (0.050) |
| log(egg area) | 0.056 (0.045) | **-0.533 (0.245)** |
| Days in pre-hatching stage | 0.001 (0.001) | **-0.004 (0.002)** |

### Egg size as predictor of other egg parameters

Species differed markedly in how well egg area predicted the other egg parameters. On average, egg area predicted egg wet weight best, then dry weight, followed by hatchling size. Overall, it predicted best for eggs of *A. elongatus* (Table 5). Models with species-specific untransformed egg area explained data better than an additive model with species and egg area effects for all the three predicted parameters (*p* < 0.001). This suggests that egg area does not predict the other parameters in the same way across different species. We then fitted egg area in separate species-specific models and calculated two complementary measures of model fit: the coefficient of determination (*R^2^*) for the fixed effects only 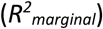 and *R^2^* of the whole model including random effects 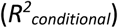. Both fixed-effects and whole-model *R^2^* for species-specific models of egg wet weight ranged widely (Table 5). Egg area was a particularly good predictor of egg wet weight in *A. elongatus* (*R^2^* = 0.959), but performed less well in the other species (Table 5). When we applied the same model to the additional *A. bellottii* dataset of wet weight from 2015, the very high value found (*R^2^_marginal_* = 0.904) contrasts with *A. bellottii* from the 2017 experiment (*R^2^_marginal_* = 0.210). In egg dry weight models, *R^2^_marginal_* was generally lower than for wet weight, again with particularly high values for *A. elongatus* from 2017 and *A. bellottii* from 2015 (Table 5). There was a small explanatory effect of egg area on hatchling size in two species that hatched in the 2017 experiment (*A. bellottii* and *A. nigripinnis*) and in *A. nigripinnis* from 2015. The exception was again *A. elongatus* from 2015 with a high *R^2^_marginal_* value (Table 5).

**Table 5.**
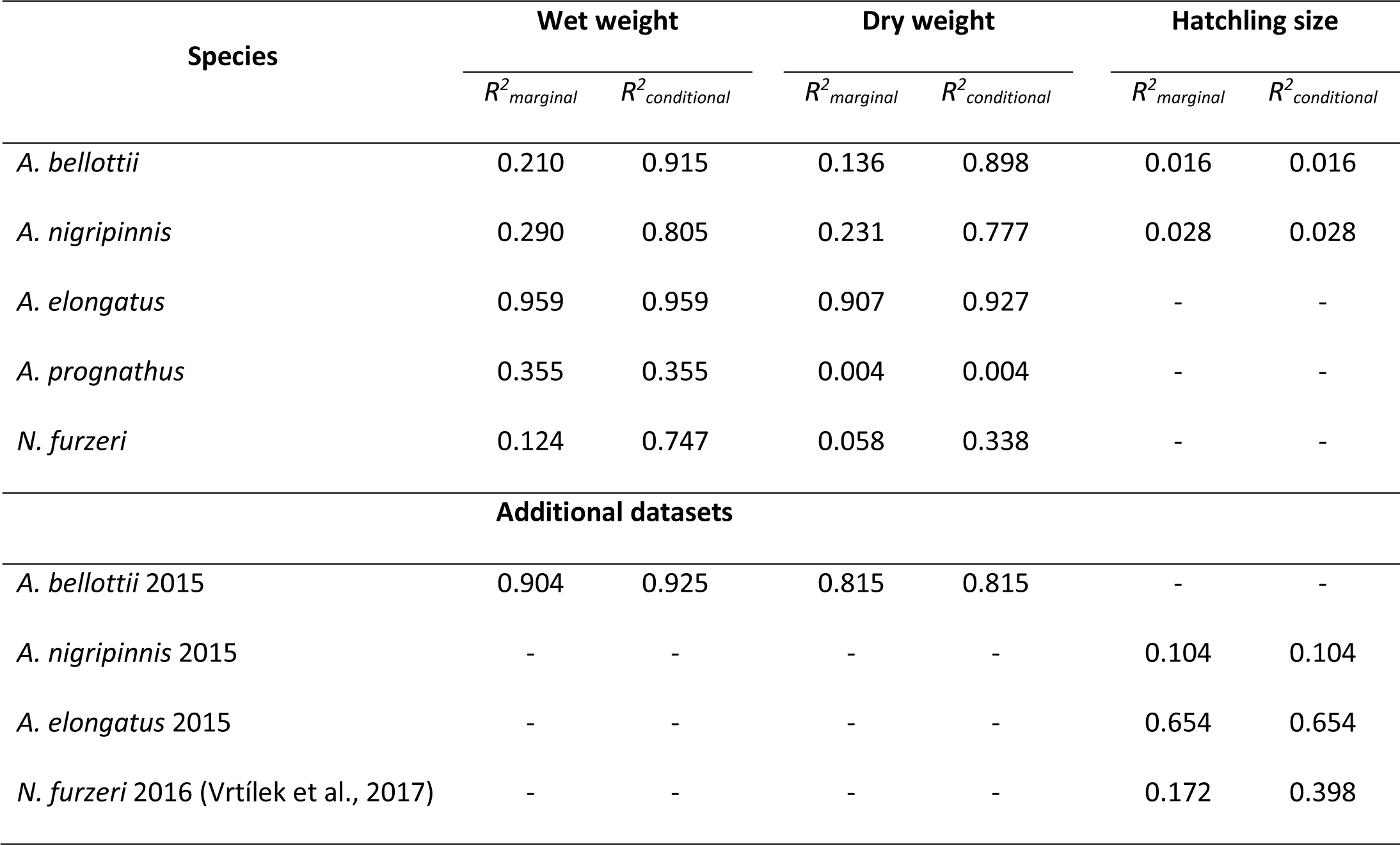
Predictive power of egg area on wet weight, dry weight and hatchling size. The *R^2^* estimate was conducted separately for fixed-effects model part (*R^2^_marginal_*) and the whole model including random effects (*R^2^_conditional_*). We compare 2017 data with the additional 2015 dataset and also with a published study on hatchling size of *N. furzeri* (Vrtílek et al., 2017).

## Discussion

Egg size can be a suitable measure of other egg parameters and offspring performance (Barneche et al., 2018; Kamler, 2005; Krist, 2011). However, can we use egg size as a universal predictor for other egg traits across species without any further adjustments? Here, we compared the composition of eggs in five annual killifish species using data obtained from individual eggs. We tested for different allometric scaling exponents of egg composition parameters on egg area. We assessed the explanatory power of egg area for variation in egg wet and dry weight and for hatchling size. We found species-specific differences in the scaling exponents of egg dry mass and water content, in elemental composition, species-specific allometries of triglyceride content and hatchling size among the studied annual killifish species. Across species, egg area performed best as a predictor for egg wet weight and within species, egg area predicted best other egg parameters in *A. elongatus,* a large piscivore species with large eggs.

### Allometric scaling exponents of egg components on egg area differ between species while patterns in exponents between traits measured on the same egg often remain similar

The analysis of allometries between traits in different species or groups is a recurring theme in morphometry (Teissier and Huxley, 1936; Klingenberg, 2016; Nakagawa et al., 2017). We can always assume equal allometries in different species, as a null model or a first working hypothesis. However, our analysis demonstrates that even within a genus, allometric scaling exponents differ between species and also within species between samples collected in different years. Even between two piscivorous species from a single clade (Loureiro et al., 2018), different scaling exponents were found. What we also observed was that traits measured on the same egg often shared the same or a similar allometric pattern. For example, egg dry mass had the same allometric exponent with egg area as water weight in each of the species except for *A. prognathus*. Amount of egg components is expected to scale isometrically with egg volume rather than with egg area and volume might have been a better measure of egg size. However, egg volume is difficult to obtain in a non-invasive manner with little handling. In the Appendix (Section B), we explain that if we assume eggs to be approximately spherical, parameter estimates of log-log regressions on egg area can be converted to the expected estimates of log-log regressions on egg volume. Slope values in log-log regressions on egg area below 1.5 imply that the response trait does not scale hyperallometrically with egg volume. For example, larger eggs of *A. bellottii* and *A. nigripinnis* had lower dry weight than expected based on a simple linear increase with egg area and also egg volume. The dry weight of *A. elongatus* eggs, however, increased more than isometrically with area but approximately isometrically with egg volume. Considering that the only slope values above 1.5 were most likely affected by outliers (in the triglyceride assay), we never found any potential hyperallometric relationship with egg volume. Parental individuals might be constrained in the amounts of resources they can allocate to each individual egg, such that they can provide at best isometric amounts.

Egg area did not affect the other egg parameters in some species. We recorded weak-to-zero relationships between egg area and dry and water weight in *A. prognathus* and *N. furzeri*. The egg area relationship with hatchling size was also absent on the intra-specific level (for both *A. bellottii* and *A. nigripinnis*). This specific finding contrasts with a previous study of *N. furzeri* with remarkably robust effect of egg size on hatchling size (Vrtílek et al., 2017) but also with *A. elongatus* 2015 data from this study. Similarly, *A. bellottii* eggs from different years showed different allometric slopes. Thus, notable sample or cohort effects are present in the analysis of annual killifish eggs, but the exact source of this variability is unknown. Moshgani and Van Dooren (2011) found that egg size and reproductive effort depended on the day when eggs were collected and attributed this to food quality and environmental state variation. Similar effects might occur for egg composition as suggested by the sample effects and the recurring presence of hypoallometries which could be caused by resource limitations.

### Water content was higher in piscivores and *N. furzeri* than in the small *Austrolebias* species

Annual killifish eggs are exposed to conditions of water stress for several months during dry season periods. Water contained in the eggs therefore represents a precious resource for the embryo. Annual killifish evolved special egg structures to prevent excessive water loss such as microvilli on the envelope or a chorion with extremely low water permeability (Machado and Podrabsky, 2007). Water content of annual killifish eggs appears on the upper boundary of the 50-70% range reported for freshwater teleosts (e.g. *Chondrostoma nasus* (Keckeis et al., 2000)). It is still much lower, however, than the 90 % recorded in marine pelagophilous fish (Craik and Harvey, 1984), where it is responsible for egg buoyancy (Lubzens et al., 2010). We found considerable interspecific variation in egg water content and the studied species formed two groups. The large piscivorous *Austrolebias* and the African *N. furzeri* had higher egg water contents than the two smaller *Austrolebias* species. This clustering is surprising from the perspective of their contrasting ecological conditions during embryogenesis. The African *N. furzeri* occurs in a subtropical semi-arid region and faces longer, colder dry periods in comparison to the temperate South-American *Austrolebias* species. In *Austrolebias*, embryos have to survive a dry season which is relatively humid but warmer. Possibly, piscivorous *Austrolebias* lay their eggs near the fringes of the temporary ponds, where these might face drought regimes more similar to the *N. furzeri* habitats. This remains an open question, however. In lab conditions, desiccation had limited effects on embryonic survival and development in *Austrolebias bellottii* (Van Dooren and Varela-Lasheras, 2018). As an alternative explanation, larger water content in eggs of some species might stem from their higher sensitivity to desiccation and not necessarily represent specialization to drier habitats.

### Elemental composition of eggs differed between species

We performed an exploratory analysis of the elemental composition of annual killifish eggs. We measured carbon content, an element which is present in various molecules acting as sources of energy; nitrogen content which provides a crude estimate of proteins; and sulphur content which is scarce in animal tissues but can be found in proteins containing methionine or cysteine amino acids (Kamler, 2005). The elemental composition of annual killifish eggs did not diverge from that of other fish such as zebrafish (46.0-54.0 % C, 9.7-11.1 % N) (Hachicho et al., 2015), Northern pike (50.7 % C, 11.5 % N) (Murry et al., 2008), or common sole (49.9 % C, 9.5 % N) (Yúfera et al., 1999). The elemental composition was specific to the different killifish species groups with the large eggs of piscivorous species containing relatively more nitrogen and sulphur when compared to the small *Austrolebias* species. In the eggs of African *N. furzeri*, we found relatively more nitrogen to carbon, again paralleling the large *Austrolebias* species.

### Egg triglycerides probably serve as an energetic reservoir for the final pre-hatching phase in annual killifish

It has been hypothesised that lipids from egg oil globules act as energy reservoir during protracted pre-hatching phase (Brind et al., 1982). In other teleost species, it is usually spent shortly after hatching (Kamler, 2008; Kohno et al., 1986; Yúfera et al., 1999). The globules contain a considerable fraction (50-60%) of the total fish egg lipids (Wiegand, 1996). In annual killifish, the proportion of total egg lipids stored in these oil globules can be particularly high, with up to 90%, and the majority of these lipids are triglycerides (Brind et al., 1982). We therefore focused our analysis on the initial amount of triglycerides in killifish eggs.

Eggs of *A. elongatus* contained larger amounts of triglycerides compared to the small species. The other piscivore, *A. prognathus*, overlapped with both *A. elongatus* and the small species. Eggs of *Z. furzeri* clustered with the small *Austrolebias* species. This suggests that the allocation of triglycerides among annual killifish species might covary with egg size. We also analysed eggs of *A. bellottii* that were incubated for over six years in damp substrate to compare their triglyceride reserves to freshly laid eggs from the same species. No embryo of these long-stored eggs had reached pre-hatching stage and there was no apparent decline of triglycerides when compared to freshly laid eggs of *A. bellottii*. Based on these findings, triglyceride levels seems to be retained until the pre-hatching state in agreement with Brind et al. (1982).

### Yolk sac reserves decrease in embryos faced with a protracted pre-hatching phase

Another energy source for fish that is mainly used during embryonic development are egg yolk lipoproteins (primarily lipovitellin) (Brooks et al., 1997; Wiegand, 1996). Freshly hatched annual killifish still possess a visible yolk sac, but an extended time of incubation (number of years) may lead to less viable hatchlings with almost spent yolk (MV, personal observation). In the two species we incubated and hatched (*A. bellottii* and *A. nigripinnis*), we monitored development regularly and, instead of hatching individual embryos at the moment when they reached the pre-hatching stage, we triggered their hatching at pre-specified date. We therefore ended up with hatchlings that had endured variable time in the pre-hatching stage. The amount of remaining yolk at hatching consequently declined with time spent in the pre-hatching stage. It seems that even during such a limited period (maximum 92 days), embryos had consumed considerable proportion of their yolk reserves. We did not inspect diapause III occurrence, so embryos may have been waiting in an active stage for a hatching cue (desiccation and inundation). This scenario may well happen in natural conditions during the last part of the dry season when humidity already increases. A larger yolk sac size would then increase persistence in the pre-hatching stage, in addition to reserves in the lipid droplet. This is similar to an increased starvation resistance during an initial post-hatching phase in other fish (Jardine and Litvak, 2003), amphibians or lizards (Moore et al., 2015; Sinervo, 1990).

## Conclusion

Egg parameter allometric scaling exponents and relationships between different egg parameters are not conserved across annual killifish species. Scaling exponents also vary between repeated samples on the same species but collected at different times in different conditions. Our results therefore call for caution and more analyses of intra- and interspecific variation in egg content allometries.

## Appendix

Analysis of allometry among species.

### A. Allometry with size

An allometric scaling describes the relationship of a trait *y* with body size *x* (or with a proxy for body size). Such relationships are usually characterized by a power law (Eqns. 1; Huxley and Teissier, 1936)

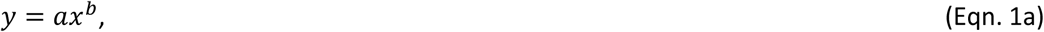

which becomes after log-log transformation

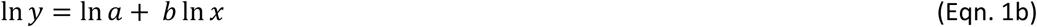

When parameter *b* (the scaling exponent) equals one then *y* is said to be isometric with body size *x*. These two traits then scale linearly by a factor *a* (the factor of proportionality). When *b > 1* we speak of positive allometry or hyperallometry; *b < 1* is called negative allometry or hypoallometry (Shingleton, 2010). A log-log regression can be used to estimate *b* and ln *a*. In its simplest least-squares or ML (maximum likelihood) formulation, the regression assumes that the residual variance around the regression line is independent of ln *x* (homoscedasticity).

### B. Different measures of body size

In regression equations used to estimate scaling exponents and factors of proportionality, size can be represented by a variable that scales with length of the individual in any life stage, the area of a section, or with its volume. In the context of this study, we assume that eggs are well approximated by considering them to be spheres with radius *r*. The area of an egg on a digital photograph then is *A = nr^2^*, whereas egg volume is 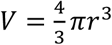. The two are related by 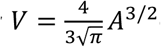 or 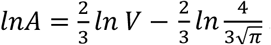 This last equation implies that the slope estimate in a log-log regression on area is two thirds the scaling exponent of an allometry with volume as a measure of size. The estimate of the factor of proportionality is also affected by this transformation and can be adjusted by adding 0.189789 to the intercept of a log-log regression on area.

### C. Allometry between traits

There are two schools in the analysis of multivariate allometries (Klingenberg, 2016). Either size corresponds to a specific measurement and can be an explanatory variable (“Gould–Mosimann” school), or it is the main axis of variation of the joint distribution of traits. This principal component is then seen as representing size variation (“Huxley–Jolicoeur” school) (Klingenberg, 2016). We follow the first approach.

When two traits *y_1_ and y_2_* both relate to body size, then we can write out their respective allometric relationships with body size *x* as

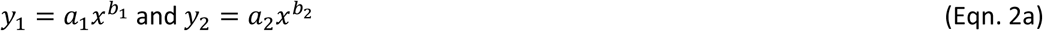

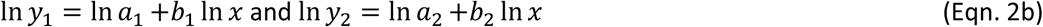

The parameters *a* and *b* can differ between traits and traits can each scale isometrically or allometrically with body size. We can also interpret the scaling between *y_1_*and *y_2_*by means of a power law:

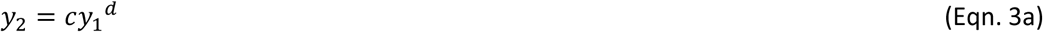

or after log-log transformation

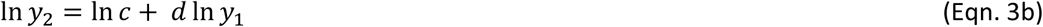

We can combine Eqns. 2 and 3 to find that

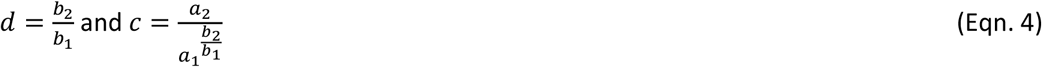

This expression parallels the one by Jolicoeur (1963), which was based on principal component analysis. Isometry with body size can differ between traits. We can also speak of isometry between a pair of traits when *d* equals one. In this case the equality *b*1 = *b*2 must hold true. To test this equality, there is no use in fitting models regressing one trait on another. Multivariate models that include both *b*1 and *b*2 as parameters provide a test, and they model the allometric relationships per trait. When both traits are isometric with body size, this equality condition is obviously satisfied as well. To test for differences in parameters *a* and *b* and to assess isometries with size and between traits, modelling all traits jointly and with body size as a covariate seems preferable, such that parameter differences can be tested by constraining equality among subsets of parameters.

### D. Regressing on body size within groups (the problem)

Following up on Gelman and Hill (2007), Nakagawa et al. (2017) discuss issues that arise when traits are regressed on body size in the presence of other covariates. Rather than being just an explanatory variable or a mediating trait, body size *x* can be a response affected by causal effects potentially shared with the other traits *y* which are regressed on it. While Shipley (2004) proposes a test to falsify potential effect orderings, (Nakagawa et al., 2017) propose not to regress on body size or log-body size as is, but to use residuals with respect to group averages for the groups defined by the other covariates modelled. This avoids that intercept terms suddenly get a different interpretation when terms including body size are removed from the model, and ensures that intercept estimates in all models compared represent average log-total amounts. These often need to be compared, while estimating factors of proportionality can be less insightful when body sizes differ much between groups. We can rewrite models for a trait *y* to accommodate this idea as

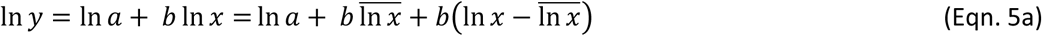

for data belonging to the first group and with *l n x* being average log-body size in that group, and

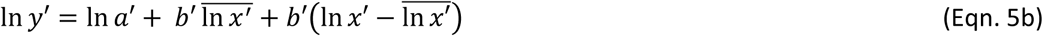

for observations *y*’ and *x*’ in the other group, for example. Log-log regressions with separate estimated intercept and slope parameters for each of these groups will estimate the allometric scaling exponents as the slopes on the group-centered log-body sizes. The factors of proportionality *ln a* and *ln a′* are not separately estimated because intercept estimates will include terms which are contributions of the average log-body size in the group scaled by the allometric scaling exponent. Intercept estimates will also be affected by conversions as explained in section B.

### E. When not to regress on centered log-body size

First of all, small sample sizes per group in the model or vastly different sample sizes can prevent fitting the centered sizes well or reduce the estimation precision on other parameters in the model. This makes it worthwhile to always fit models without regressions on body size as well, to check the stability of estimates of other parameters. In any case where we can’t include sizes or centered sizes in models, we need to remain aware that the interpretation of an intercept becomes the one based on Eqns. 5: it is the factor of proportionality of an allometric relationship plus a term depending on the average size scaled by the allometric exponent.

A second issue arises when a response variable *y* is calculated directly from body size, i.e. when the calculation of the response involves body size or its proxy. This becomes a regression of a variable on itself, amply discussed and criticised by Nee et al. (2005). We then need to resort to models that don’t have explicit body size terms, hence intercepts again include a term depending on average size scaled by the allometric exponent.

A third issue arises in the context of analysing concentrations or compositions. An analysis of a response which is a ratio of a trait *y* and *x* by means of a model which includes a body size covariate, can in fact be interpreted as a model that fits a body size term to test whether the dependence between *y* and *x* deviates from isometry. A side effect of analysing a ratio is that it becomes less clear what the pattern of residual variance in the ratio should be. A useful option seems to be using a log-log regression including log-body size as an offset term (i.e. specifying log-body size coefficient as 1 and thus assuming isometry). Then the response modelled is the trait *y*, for which variance properties are better known or less constrained. In the case where we assume isometry (we analyse the concentration *y*/*x*), there is no parameter *b* estimated. When data don’t allow regressing on body size terms to test for deviations from isometry, we can interpret all fitted terms as if we assume isometry but it seems better to say that estimated intercept terms include a term depending on average size scaled by the allometric exponent minus one. For the analysis of compositions, dedicated methods exist (van den Boogaart and Tolosana-Delgado, 2013). If model parameters are not easily interpretable, we can switch to a multivariate analysis of total amounts to interpret parameters, while testing contributions of covariates can be done according methods for compositional data. This approach avoids the use of ratios, but a proper model for compositional data is replaced with a model compositional methods were invented to avoid.

### F. Measurement error

Warton et al. (2006) discuss at length the consequences of measurement and equation error in the analysis of bivariate allometries. They state that, in many instances, the two traits involved are equally important and that therefore major axis (MA) or reduced major axis (RMA) are obvious least-squares model-fitting choices. In a likelihood approach, one would then resort to likelihoods based on multivariate distributions for each observation and the slope *b* becomes part of the specification of the covariance between traits (Warton et al., 2006). Covariance regression models can then be used to test for differences in slope *b* between groups. Voje et al. (2014) on the other hand, do opt for a regression approach where body size is seen as an explanatory variable. In the models we fitted to several traits, such a regression approach was implemented and no covariance regressions were fitted for simplicity. When the main goal of inference is predicting a trait on the basis of body size, standard regression models should be used (Warton et al., 2006). These are also most adequate in the absence of measurement error in body size or they suffice when the goal of inference is to test whether body size significantly affects another trait, i.e. whether *b* is non-zero or different from one in the case of concentrations in models with an offset. In the presence of measurement error estimates of regression slopes are affected, therefore when we need reliable estimates of *b*, measurement error needs to be considered and taken into account. However, across a number studies inspected by Warton et al. (2006) the effects seemed limited to a decrease in the estimated value of the slope *b* which is below 8%. Next to other methods such as the method-of-moments regression (Warton et al., 2006), Carroll et al. (2006) propose regression calibration and simex extrapolation as generally applicable methods to derive regression slopes corrected for measurement error in body size which still permit standard regression or mixed models can be fitted and the associated inference methods used.

### G. Explaining variation in traits dependent on body size

To calculate coefficients of determination or generalized coefficients of determination, regression models are required that regress a response trait *y* on body size *x*. Here, we motivate our choice to do this in linear regressions of *y* on *x* and not by means of log-log regressions. In log-log regressions, terms including body size estimate allometric exponents. So, if we calculate coefficients of determination for these regression terms, we don’t assess the contribution of the parameters *a*, which are the primary scaling parameters we are interested in as a description of the explanatory effects. Here, we are not interested in the allometric exponents that control curvature of the allometry and we would also need to interpret coefficients of determination for intercepts, which is counterintuitive. We therefore analyse response of raw *y* on size *x* values. The non-linear effects then contribute to unexplained variation of the linear regression.

## Acknowledgments

We would like to thank Simon Agostini (CEREEP) and Jakub Žák (IVB, CAS) for their help with fish husbandry. We are thankful to Jakub Hruška and Milena Žídková (Global Change Research Institute, CAS) for measuring the elemental composition of eggs. We also want to thank Véronique Vaury (IEES, Sorbonne University) for helping with egg weighing, and Sandrine Meylan (IEES, Sorbonne University) for discussion on the triglyceride content analysis. We thank Radim Blažek (IVB, CAS) for photographs of live fish.

Funding: MV was supported by French government scholarship program (Bourse du Gouvernement Français Stage 2017). This study was funded by Pepinière Evo-Devo PSL and Czech Science Foundation project 19-01781S.

## Supplementary Information

Supplementary information for this article is available at publisher’s website.

Data and R script accompanying this article are available under DOI 10.6084/m9.figshare.11897247

