## Supplemental Table 1 for "Egg size does not universally predict embryonic resources and hatchling size across annual killifish species"

Table S1. Age and total length of the spawning pairs in the 2017 experiment.

Table S1. **Age and total length of the spawning pairs in the 2017 experiment.** Age of females is given in weeks when the body size was measured (on August 7, 2017). Species codes: AB – *Austrolebias bellottii* “Ingeniero Maschwitz” (small), AN – *A. nigripinnis* “La Guarderia” (small), AE – *A. elongatus* “General Conesa” (piscivore), AP – *A. prognathus* “Salamanca” (piscivore), NF – *Nothobranchius furzeri* “MZCS414” (small - African).

| **Tank** | **Species** | **Female age** (Hatch date – dd/mm/yyyy) | **Male TL** | **Female TL** | **Spawned/ Incubated eggs** |
|---|---|---|---|---|---|
| 4 | AP | 28 (26/01/2017) | 79 | 75 | 26/13 |
| 5 | AE | 61 (08/06/2016) | 85 | 68 | 35/9 |
| 7 | AE | 61 | 87 | 78 | 55/17 |
| 8 | AP | 28 | 86 | 97 | 64/25 |
| 15 | AN | 75 (04/03/2016) | 40 | 37 | 601/66 |
| 16 | AB | 28 | 69 | 48 | 928/397 |
| 17 | AN | 28 | 50 | 43 | 1433/261 |
| 18 | AB | 28 | 63 | 54 | 1450/108 |
| 19 | AN | 75 | 48 | 43 | 588/77 |
| 20 | AB | 41 (26/10/2016) | 56 | 55 | 729/68 |
| 21 | AN | 28 | 41 | 39 | 1199/11 |
| 22 | AB | 28 | 59 | 53 | 1289/256 |
| 102 | AE | 36 (28/11/2016) | 77 | 60 | 79/47 |
| T1 | AE | 41 (26/10/2016) | 70 | 65 | 48/14 |
| T2 | AE | 41 | 60 | 59 | 17/15 |
| N | NF | 5 (4/7/2017) | 36 | 37 | 22/- |
| N | NF | 5 | 49 | 36 | 17/- |
| N | NF | 5 | 45 | 37 | 23/- |
| N | NF | 5 | 52 | 44 | 19/- |
| N | NF | 5 | 35 | 35 | 34/- |
| N | NF | 5 | 37 | 36 | 2/- |
